# Deep learning diffusion fingerprinting to detect brain tumour response to chemotherapy

**DOI:** 10.1101/193730

**Authors:** Tom A. Roberts, Ben Hipwell, Giulia Agliardi, Angela d’Esposito, Valerie Taylor, Mark F. Lythgoe, Simon Walker-Samuel

## Abstract

Artificial neural networks are being widely implemented for a range of different biomedical imaging applications.Convolutional neural networks are by far the most popular type of deep earning architecture,but often require very large datasets for robust training and evaluation We introduce deep learning diffusion fingerprinting (DLDF), which we have used to classify diffusion-weighted magnetic resonance imaging voxels in a mouse model of glioblastoma (GL261 cell line), both prior to and in response to Temozolomide (TMZ) chemotherapy.We show that, even with limited training, DLDF can automatically segment brain tumours from normal brain, can automatically distinguish between young and older (after 9 days of growth) tumours and that DLDF can detect whether or not a tumour has been treated with chemotherapy.Our results also suggest that DLDF can detect localised changes in the underlying tumour microstructure, which are not evident using conventional measurements of the apparent diffusion coefficient (ADC).Tissue category maps generated by DLDF showed regions containing a mixture of normal brain and tumour cells, and in some cases evidence of tumour invasion across the corpus callosum, which were broadly consistent with histology.In conclusion, DLDF shows the potential for applying deep learning on a pixel-wise level,which reduces the need for vast training datasets and could easily be applied to other multi-dimensional imaging acquisitions

**Abbreviations:**

ANN: artificial neural network
CT: x-ray computed tomography
PET: positron emission tomography
CNN: convolutional neural network
HARDI: high angular resolution diffusion weighted imaging
NODDI: neurite orientation dispersion and density imaging
VERDICT: vascular, extracellular and restricted diffusion for cytometry in tumours
DLDF: deep learning with diffusion fingerprinting
TMZ: Temozolomide
PFA: paraformaldehyde
H&E: hematoxylin and eosin
GFAP: glial fibrillary acidic protein

## 1. INTRODUCTION

Research into the use of deep learning with artificial neural networks (ANN) is being widely undertaken.A key application is in radiology, particularly in the classification and segmentation of biomedical images (1–4).ANNs have been trained and evaluated using x-ray computed tomography (CT) (5–7) and positron emission tomography (PET) imaging (8–10), and several magnetic resonance imaging (MRI) data types including structural T1- and T2-weighted images (11), perfusion images (12), MR spectroscopy (13) and diffusion MRI (14).

In the last five years,convolutional neural networks (CNNs) have become by far the most popular choice of deep learning architecture (15).CNNs allow subtle and abstract features to be characterised, but often require very large, labelled data sets (16).As an alternative approach, we propose using data-rich diffusion-weighted MRI acquisitions to define a fingerprint in each voxel, which can easily be classified according to tissue type and used to traina deep neural network.This approach aims to produce accurate diagnostic information.with a relatively small number of subjects,as every voxel is treated as an independent data point to train the ANN,rather than using whole images (or patches from within images) as used in CNN architectures.

Contrast in diffusion weighted MRI is dependent on the random motion of water molecules and the structure of the underlying tissue microenvironment.Various multi-direction and multi-b-value diffusion MRI protocols have been proposed which are better at resolving complicated tissue microstructure compared to basic diffusion tensor imaging(DTI). HARDIhigh angular resolution diffusion weighted imaging) and NODDI (neurite orientation dispersion and density imaging) use multiple diffusion directions (17,18) to improve measurement of neuronal fibre direction, whilst VERDICT (vascular, extracellular and restricted diffusion for cytometry in tumours) protocols employ multiple directions and multiple sets of diffusion weighted scans (19, 20) to make estimates of cancer cell microstructure and density.These types of diffusion MRI protocols are particularly amenableto pixel-wise deep learning because large amounts of data are acquired inindividual subjects.

A key advantage of deep learning over techniques which require mathematical models to fit the acquired diffusion signals, is that no assumptions are made about the underlying tissue microstructure.This is especially important when imaging pathology where, often, the lesiongreatly alters or interferes with the tissue microstructure in the affected organ.Since most mathematical models are optimised to characterise one type of tissue, regions of pathology can often be poorly characterised because they simply correspond to regions where themodel fitting is less robust.For example, NODDI is not designed to characterise brain tumour microstructure and VERDICT is not designed to characterise normal brain tissue. Characterising microstructure in transitional regions containing both neuronal and cancercells is even more challenging.

This therefore underpins our motivation for developing deep learning with diffusion fingerprinting (DLDF).In this study, we have evaluated DLDF in a set of diffusion MRI data from a mouse glioma model (GL261), both prior to and following treatment withTemozolomide (TMZ) chemotherapy. We investigate the ability of DLDF to distinguish brain, background, control and treated tumour pixels, independent of pixel location, and without applying mathematical models of tissue microstructure.

## 2. METHODS

### 2.1 Ethics statement

All in vivo experiments were performed in accordance with the UK Home Office Animals Scientific Procedures Act, 1986 and United Kingdom Coordinating Committee on Cancer Research (UKCCCR) guidelines, and with the approval of the University College London Animal Ethics Committee.

### 2.2 Mouse Glioma Model and Therapy

Female, C57BL/6 nude mice (6-9 weeks old) were sourced from Harlan (UK) and were housed in groups of 5 in isolated ventilated cages (IVCs) containing environmental enrichment.Mice had access to food and water ad libitum.

Mice were injected in the right striatum with 2x10^4^ GL261 mouse glioma cells, and randomly assigned to control and treatment (Temozolomide (TMZ)) groups (n = 12 per group) Following 13 days of tumour growth (day 0), to an average tumour volume of 8.1 ± 0.8 mm^3^,three doses of TMZ (130mg/kg) or vehicle were administered by gavage, on consecutive days.

### 2.3 MRI setup

MRI measurements were performed on a 9.4T horizontal bore scanner (Agilent Technologies, Santa Clara, USA) with a 205/120/HD 600mT/m gradient insert. RF transmission was performed with a 72mm inner diameter volume coil and a 2-channel receiver coil (RAPID biomed, Ripmar, Germany).Mice were anaesthetised with 1.5-2.0% isoflurane in 2 l/min oxygen and positioned prone in a cradle for imaging. The head was positioned within an MR-compatible head-holder and secured with plastic ear bars to minimise motion artefacts. Body temperature was maintained at physiological temperature using a hot water system and monitored using a rectal probe (SA Instruments).Respiration was monitored using a neonatal apnoea pad taped to the abdomen of the animal.For gadolinium-based scans,an intraperitoneal infusion line was inserted into the mouse.

### 2.4 Diffusion MRI

Gliomas were localised using a structural spin-echo sequence. For generating the diffusion fingerprints, diffusion-weighted images were acquired in a coronal orientation using a 3-shot spin-echo echo planar imaging (EPI) sequence, with the following parameters: TR = 3s, TE= min, DM = 64^2^, FOV = 20mm^2^, shots = 3, slice thickness = 0.5mm, slices = 5, averages =2. In total, 46 diffusion weightings in 3 directions were acquired in addition to a 42 direction DTI acquisition (b = 1000 s/mm^2^). Specific gradient combinations are detailed in Table 1. TEwas minimised for all scans to maximise signal-to-noise.To correct for signal changes caused by this variation in TE, an accompanying b = 0 s/mm2 (B0) image was acquired for every combination of diffusion gradients.Total acquisition time for all DWIs was 70 minutesFollowing diffusion imaging, mice were injected with gadolinium-DTPA (Magnevist). After 10minutes – to allow for the contrast agent to circulate – slice-matched T1-weighted spin echo EPI images were acquired. Tumour regions of interest (ROI) were drawn based on these images and applied to the diffusion data.

**Table 1:** Diffusion gradient combinations used in mouse brain imaging for creating diffusion Fingerprints.

| $\delta$ (ms) | $\Delta$ (ms) | G (G/cm) | b-value (s/mm<sup>2</sup>) |
| --- | --- | --- | --- |
| 3 | 10 / 20 / 30 / 40 | 3.6 | 8 / 16 / 24 / 33 |
| 3 | 10 / 20 / 30 / 40 | 7.2 | 30 / 63 / 97 / 130 |
| 3 | 10 / 20 / 30 / 40 | 10.8 | 68 / 143 / 218 / 293 |
| 3 | 10 / 20 / 30 / 40 | 14.4 | 120 / 254 / 387 / 521 |
| 3 | 10 / 20 / 30 / 40 | 18.0 | 188 / 397 / 606 / 814 |
| 3 | 10 / 20 / 30 / 40 | 21.6 | 270 / 571 / 872 / 1173 |
| 3 | 10 / 20 / 30 / 40 | 25.2 | 368 / 778 / 1187 / 1596 |
| 3 | 10 / 20 / 30 / 40 | 28.8 | 481 / 1016 / 1550 / 2085 |
| 3 | 10 / 20 / 30 / 40 | 32.4 | 609 / 1285 / 1962 / 2639 |
| 3 | 10 / 20 / 30 / 40 | 36.0 | 752 / 1587 / 2422 / 3257 |
| 10 | 30 / 40 | 4.0 | 305 / 420 |
| 10 | 30 / 40 | 8.0 | 1221 / 1680 |
| 10 | 30 / 40 | 12.0 | 2749 / 3780 |

### 2.5 Histology

To compare the DLDF category maps with a ground truth, histology was performed on the mouse brains.After the final MRI scan, mice were terminally anaesthetised via intraperitoneal injection of 100 mg/kg sodium pentobarbital (Pentoject, Animalcare, York,UK) and then transcardially perfusion-fixed with 10 ml heparinised saline followed by 10 ml 4% paraformaldehyde (PFA). For histology, mouse brains were extracted, sliced in a coronal orientation and stained with hematoxylin and eosin (H&E) and glial fibrillary acidic protein(GFAP).

### 2.6 Labelling and Diffusion Fingerprint Preparation

Four tissue categories were manually identified on contrast-enhanced images: 1)background (corresponding to any pixel that was not part of any other region), 2) normal-appearing brain, 3) untreated tumour (regions showing contrast-enhancement in all animals at day 0 and only control animals at day 9) and 4) treated tumour at day 9 (Figure 1a) Diffusion fingerprints consisting of 232 ordered data points, from individual pixels (Figure 2) were produced for each pixel, and were normalised to an acquisition with no diffusion Encoding.

### 2.7 Deep Learning of Diffusion Fingerprints

We created a deep neural network with 5 hidden layers in Keras (a model-level library, Python 3.5.2) using TensorFlow as the numerical backend (21). Each hidden layer was regularized with 20% dropout (to avoid overfitting), and used rectified linear unit (ReLU) activation and normally-distributed initialisation.The number of nodes for each layer was 100, 150, 200, 512 and 1024, respectively. The output layer contained 6 nodes corresponding to the categories defined above.Diffusion fingerprints from individual pixels were passed to the network input layer.

### 2.8 Training

50% of the data were randomly assigned to a training set, which were further refined to include equal numbers of fingerprints from each category (a total of 6,608 fingerprints) Training of the network was performed on a Nvidia Titan-X GPU, using stochastic gradient-descent optimisation (learning rate, 0.01; decay, 1e-6; momentum, 0.9), with mean squared error loss function Training was performed for 50,000 epochs, with 10,000 steps in each.

### 2.9 Evaluation

The evaluation data set contained 5 animals per group. Each diffusion fingerprint was categorised by the deep learning network and reconstructed as category maps. Dropout regularisation was used to estimate classification probability and variance via Monte Carlo sampling (22).

## 3. RESULTS

### 3.1 Brain Region Classifiers and Diffusion Fingerprint Generation

Gliomas were hyperintense on T1-weighted post-gadolinium images compared to normal brain at both day 0 and day 9 (Figure 1a).These images were used for manual delineation of brain and tumour regions (Figure 1a, dashed lines) resulting in four categories (Figure 1b):outside of brain, ‘normal’ brain, untreated tumour (all day 0 tumours and day 9 tumours which received no treatment) and treated tumour (day 9 tumours, which were treated with Temozolomide).

**Figure 1:**
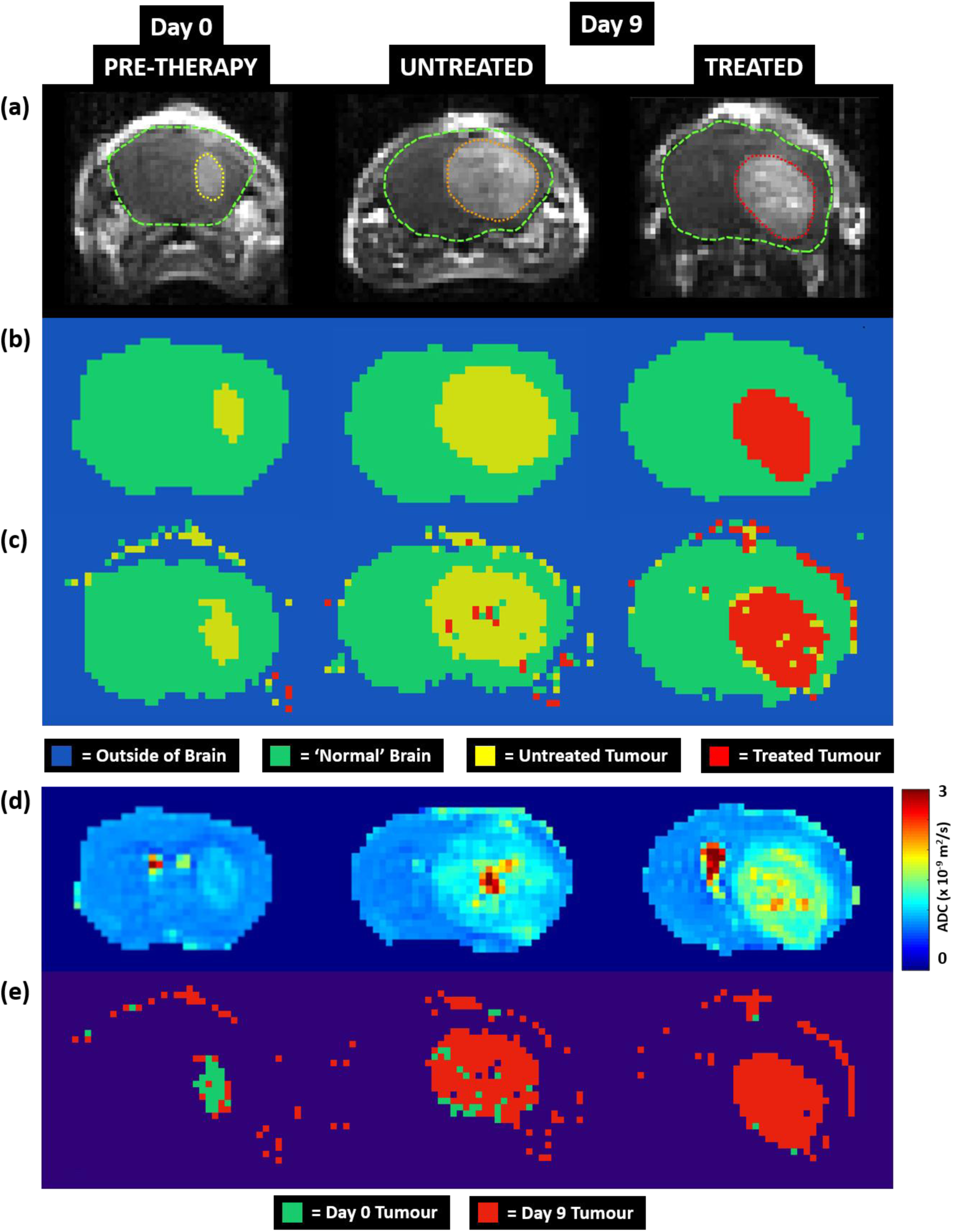
Coronal images from mouse brains at day 0 and day 9. (a) T1-weighted post-gadolinium slices showing manual segmentation (dashed lines).(b) Masks showing classification regions used for training the deep learning neural network.(c) DLDF category maps from the evaluation phase.(d) ADC maps.(e) Maps of tumour age from the evaluation Phase.

The category masks were applied to the slice-matched diffusion data to generate the diffusion fingerprints required for training the ANN. Figure 2a shows a complete dataset of diffusion MR images from one coronal slice through a mouse brain.Image intensity decreases as the level of diffusion-weighting is increased.Figure 2c shows diffusion fingerprints from four different regions.Diffusion fingerprint 1 (blue) comes from a region outside of the animal, hence the plot is a noisy trace.Diffusion fingerprint 2 (green) comes from normal brain, whilst diffusion fingerprints 3 (yellow) and 4 (red) originate from untreated and treated tumour, respectively.The first 48 points in each plot represent the DTI (plus six B0) scans. The remaining points represent different combinations of diffusion gradients (detailed in Table 1).Upon visual inspection, some differences can be observed in the diffusion fingerprint from the normal brain compared to the fingerprints from the tumours.For example, S/S0 for the DTI scans is ∼0.5, compared to ∼0.3–0.4 in the tumours.Furthermore, S/S0 decays more quickly in the tumours. The differences between the untreated (yellow) and treated tumour (red) fingerprints are subtler, however, demonstrating the need for deep learning to tease out the features within these fingerprints.

**Figure 2:**
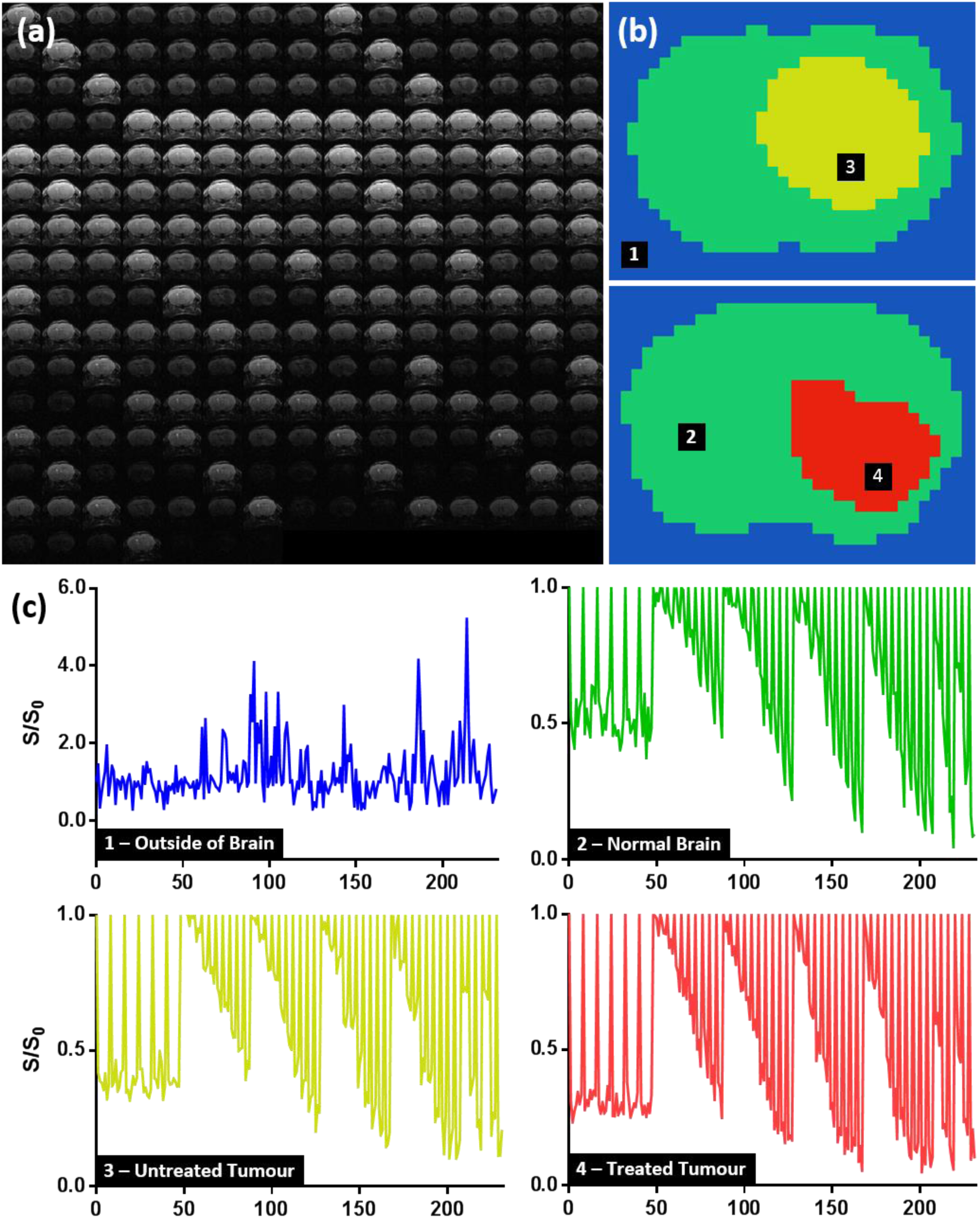
(a) Complete set of diffusion MR images from a single coronal slice in the brain. (b) Classification masks from an Untreated tumour and a Treated tumour.(c) Example diffusion fingerprints taken from the corresponding numbered region in panel (b), showing the normalised diffusion signal taken from a single voxel.

### 3.2 Model Accuracy

In total, 6,608 fingerprints from 50% of the animals were used to train the ANN. Training lasted 80 hours, and resulted in a model accuracy of 98.65% (where pixels were classified as the same category determined by manual segmentation).Based on evaluation of the ANN using the remaining 50% of the imaging data, accuracy was 98.54% (once again, compared to manually segmented data).

#### 3.3 Voxel Classification

A strong agreement was observed between manually segmented category maps (Figure 1b) and DLDF maps generated from the evaluation phase (Figure 1c) demonstrating the ability of the ANN to automatically segment tumour from normal brain.Predicted tumour volumes were 2.1 ± 0.2 times larger than manually-segmented volumes.This is due in part to the identification of tumour deposits that had grown beyond the skull.Ghosting artefact in background regions was also miscategorised as brain or tumour tissue.

The ANN was able to reliably identify diffusion fingerprints corresponding to normal brain (green),untreated tumour (yellow) or post-treatment tumour (red).Furthermore, the ANN was able to distinguish between tumours based on age (Figure 1e): voxels within day 0tumours were predominantly classified as day 0 pixels (green, 92% accuracy), whilst untreated and treated tumours were mostly classified as day 9 pixels (red, 93% accuracy) Regardless of age, tissue outside of tumours was predominantly classified as normal brain (purple, 97% accuracy).

In comparison, ADC alone was unable to correctly stratify all pre-therapy tumours from untreated tumours by simple thresholding (Figure 3b).Whilst the mean ADC value in the pre-therapy tumours was significantly lower than the untreated tumours at day 9 (p < 0.001), several tumours in both groups exhibited ADC values between 1.05 x 10^-9^ and 1.20 x 10^-9^ m^2^/s (blue band, Figure 3b).The mean ADC value in the treated group at day 9 was 47% higher than baseline and all mice could be distinguished from pre-therapy and control tumours.

**Figure 3:**
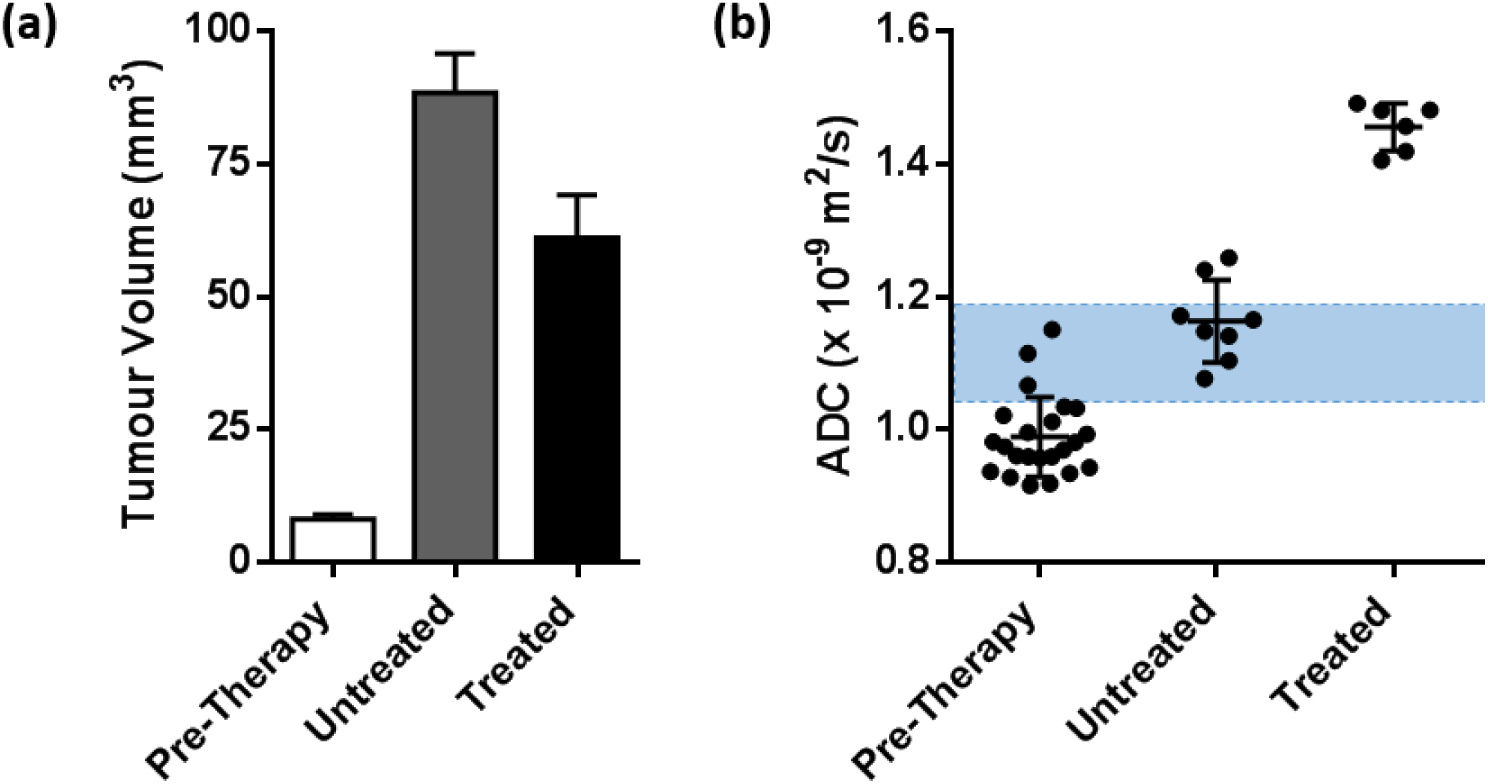
(a) Tumour volume measured from T2-weighted images. (b) Mean ADC measurements from mouse brain tumours (p < 0.001, Kruskal-Wallis one-way ANOVA).The blue region denotes a range of ADC values where pre-therapy and untreated tumours cannot be stratified based on ADC alone.

**Figure 4:**
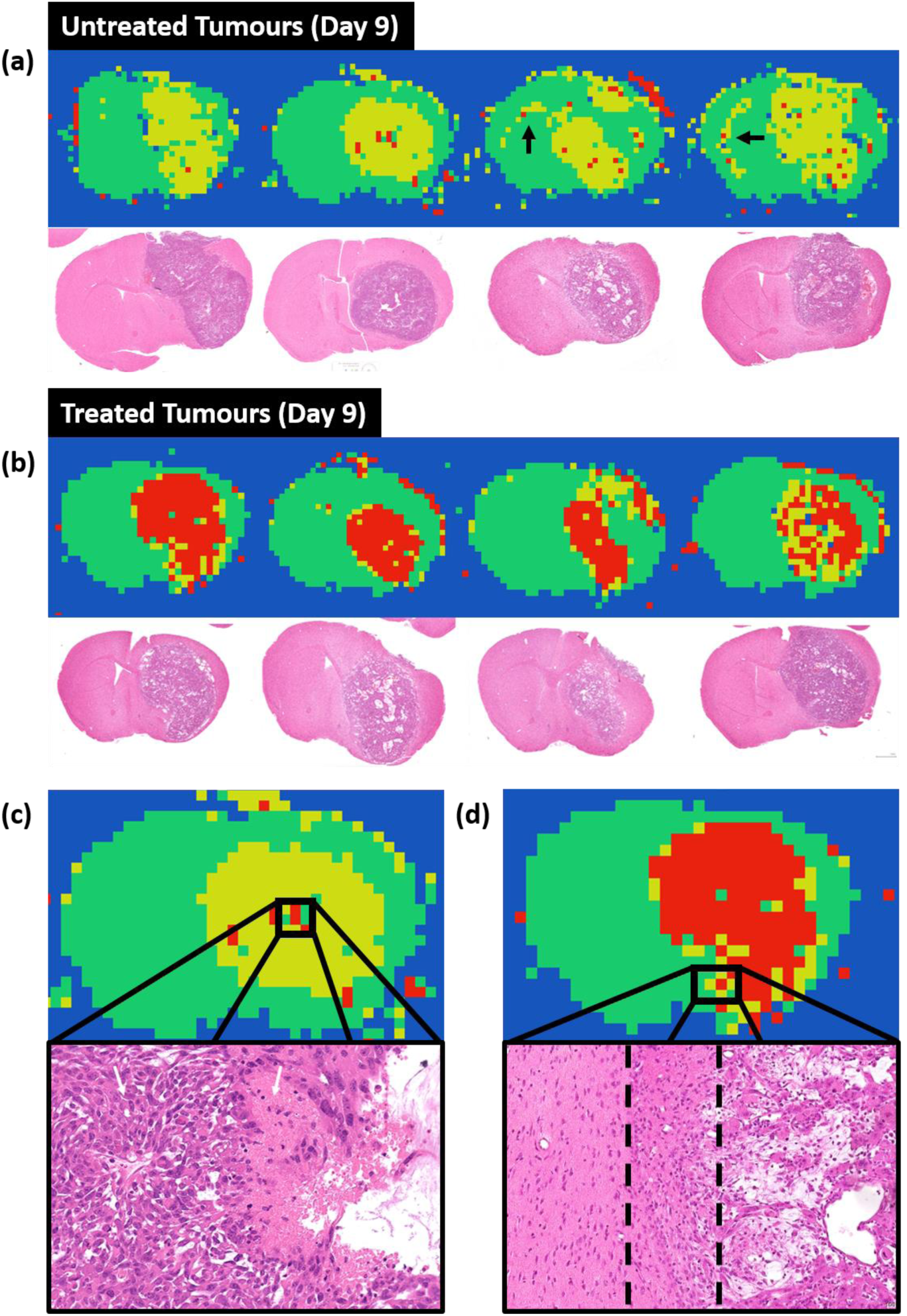
DLDF classification maps and corresponding H&E histological sections at day 9 in Untreated tumours and (b) treated tumours. DLDF could correctly distinguish between all untreated and treated tumours at Day 9.(c) and (d) Localised regions in DLDF classification maps and the corresponding regions on histology.The untreated tumour in (c) shows voxels classified as normal brain, which correspond approximately with neuronal tissue on histology The treated tumour in (d) shows a transition in voxel classification (left to right) from normal brain (green) to untreated tumour (yellow) to treated tumour (red), which corresponds to three distinct regions on histology: normal brain parenchyma, a region of tumour infiltration and the bulk tumour tissue.

Visual inspection of DLDF category maps from untreated tumours at day 9 (Figure 4a) revealed a more spiculated appearance than treated tumours at day 9 (Figure 4b) and showed a strong spatial accordance with H&E-stained histological sections.Centralised regions of normal brain pixels (green) and ‘treated’ pixels (red) were often observed inside the untreated tumours. These regions could correspond to areas on histology (Figure 4c) which showed a mix of tumour cells, neuronal tissue and voids likely corresponding to cell death.

DLDF category maps from treated animals generally showed a distinctive rim of ‘untreated’ pixels (yellow) at the tumour periphery (Figure 4b), which could correspond to border regions in H&E images where the tumour transitions into the normal brain (Figure 4d).Whether these are regions that truly show less sensitivity to TMZ therapy requires further investigation.

**Figure 5:**
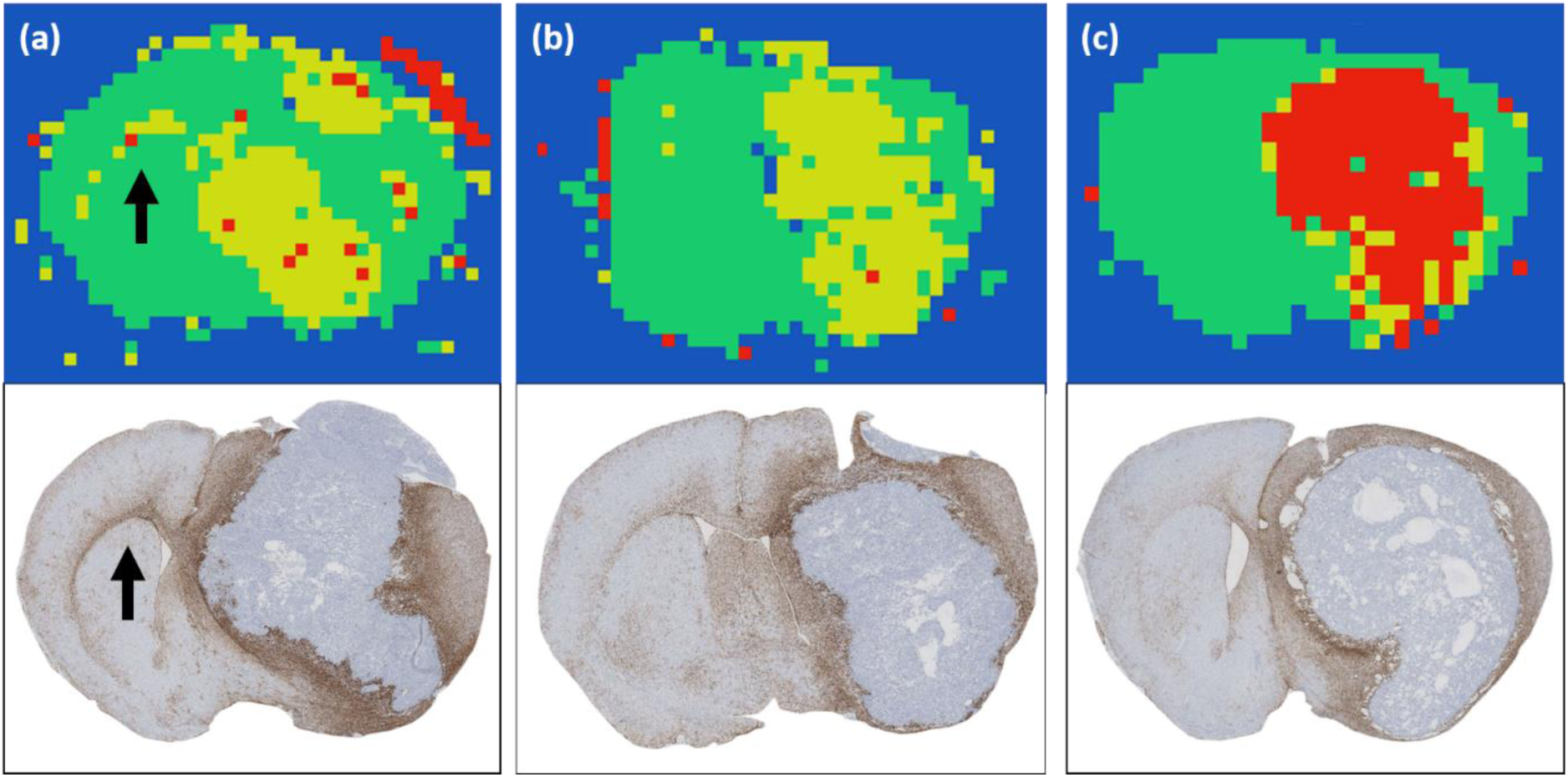
DLDF category maps compared with GFAP-strained histological sections from the same brains.(a) Evidence of tumour invasion across the corpus callosum in the DLDF map of an untreated tumour corresponds to a region of enhanced GFAP staining (black arrows).DLDF maps of (b) other untreated and (c) treated tumours showed no indication of corpus callosum invasion, which was reflected in histology.

In some of the untreated animals (Figure 5a, black arrows), the predicted category maps showed tumour invasion into the contralateral hemisphere along the corpus callosum Histological sections from the same brains showed increased GFAP-staining along the corpus callosum (Figure 5a), whereas much less GFAP-staining was visible in untreated (Figure 5b) and treated (Figure 5c) mouse brains, which displayed no evidence of tumour invasion on DLDF category maps.

### 4. DISCUSSION

The aim of this study was to evaluate the ability of DLDF in a mouse brain tumour model to categorise normal brain, pre-treatment, untreated and Temozolomide-treated tumour pixels. We have shown that DLDF can automatically segment tumours from normal brain, DLDF can automatically distinguish between young and older (after 9 days of growth) tumours and that DLDF can detect whether or not a tumour has been treated with chemotherapy We have also shown that, spatially, the category maps generated by DLDF were broadly consistent with histology and that interesting features are detected which may correspond to transitional regions within tumours where the microstructure is a mixture of normal brain and tumour cells.

The ANN used for DLDF was trained on only 6,608 fingerprints and achieved good prediction and evaluation accuracies.Broadly,the shape of the tumours delineated using DLDF were consistent with manually-segmented training data. Interestingly, during manual segmentation of T1-weighted images, regions of enhancement above the bulk of the tumour or even external to the skull, which were likely caused by deposition of tumour cells during retraction of the inoculation needle, were not segmented as tumour, and yet these regions were correctly labelled as tumour on DLDF category maps.

Likewise, we also found evidence in DLDF category maps for tumour invasion into the contralateral hemisphere, via the corpus callosum, but only in control mice.This was accompanied by a much more spiculated appearing boundary, which contrasted with the smoother appearance of TMZ-treated tumours.No evidence of invasion was detected in treated mouse brains, most likely as a direct result of TMZ causing growth inhibition. GFAP- stained histological slices from the same brains confirmed these regions of invasion,where more intense staining was seen along the corpus callosum in the untreated animals.This finding is of particular interest, as these invasion effects were not included in the manually segmented ROIs used to train the ANN. It also presents a challenge in regard to quantifying the accuracy of the ANN, as these, and other, correctly segmented and informative regions actually serve to reduce the reported accuracy of the ANN.

Further deviations between manual segmentation and the DLDF category maps were evident,and which were consistent with histology.Untreated tumours at day 9 contained numerous voxels that were categorised as normal brain,and which were positioned in a similar location to regions of brain tissue within the tumours. Likewise,voxels at the periphery of TMZ-treated tumours were often misclassified as ‘untreated’. This could be a straightforward misclassification error by the ANN, or evidence of regions of tumour that were less prone to changes in microstructure induced by Temozolomide, or even with a potential for resistance. This could be an interesting area for further research, as\ noninvasive biomarkers of treatment resistance would be particularly useful in the clinic for monitoring response (23,24).

The DLDF results are also interesting as they appear to outperform an alternative, more conventional approach to treatment monitoring, which is to apply simple thresholding to mean tumour ADC measurements. Untreated brain tumours could be distinguished from treated tumours based on their mean ADC, which was significantly higher in treated mice. However, assessment of localised treatment response was not possible using ADC alone. In treated mice, ADC was broadly elevated across the whole extent of tumour, however, untreated mice exhibited localised regions of similarly elevated ADC, most likely corresponding to regions of necrosis or (non-chemotherapy related) apoptosis. DLDF provides a localised map of treatment response, possibly highlighting regions of non-responding, or chemotherapy resistant, tumour tissue.

The results presented in this study, although preliminary, show some of the potential for deep learning on an individual voxel level, and could be applied to other types of multi- dimensional acquisitions. Even with limited training DLDF was able to accurately classify normal brain, treated tumours and control tumours at two different time points, and in some regards, was able to out-perform manual segmentation by identifying regions of tumour invasion, mixed tissue types and, potentially, tumour regions that were less sensitive toTMZ-treatment. The ability of DLDF to predict response to therapy and to ‘age’ tumour pixels will be the focus of further investigation.

## Acknowledgments

Wellcome Trust Senior Research Fellowship (grant WT100247MA), KCL and UCL Comprehensive Cancer Imaging Centre CR-UK & EPSRC (In association with the DoH England), the EPSRC-funded UCL Centre for Doctoral Training in Medical Imaging (EP/L016478/1).

